# Creating Coarse-Grained Systems with COBY: Towards Higher Accuracy in Membrane Complexity

**DOI:** 10.1101/2024.07.23.604601

**Authors:** Mikkel D. Andreasen, Paulo C. T. Souza, Birgit Schiøtt, Lorena Zuzic

## Abstract

Current trends in molecular modelling are geared towards increasingly realistic representations of the modelled systems. This is reflected in larger, more complex systems, which are difficult to build and would ideally rely on a software that converts user-provided descriptors into system coordinates. This is not a trivial task, as the building algorithms use simplifications that can introduce inaccuracies in the system properties that do not correspond to the requested values. We created COBY, a coarse-grained system builder that can create a large variety of systems in a single command call using Martini molecule models. We improved the accuracy of the complex membrane and solvent building procedures, introduced a variety of arguments that can be used to build diverse systems, and implemented features intended for force field development. COBY can be used to build flat membranes of any degree of complexity, handle protein and solvent insertion, solute flooding, stacked membranes, membrane patches and pores, and includes advanced functionalities such as molecule import, lipid building from fragments, handling of multiple parameter libraries, and several choices of algorithms for interpreting user-provided system descriptors. COBY is an open-source software written in Python 3, and the code, documentation, and tutorials are hosted at github.com/MikkelDA/COBY.

**TOC Graphic:** 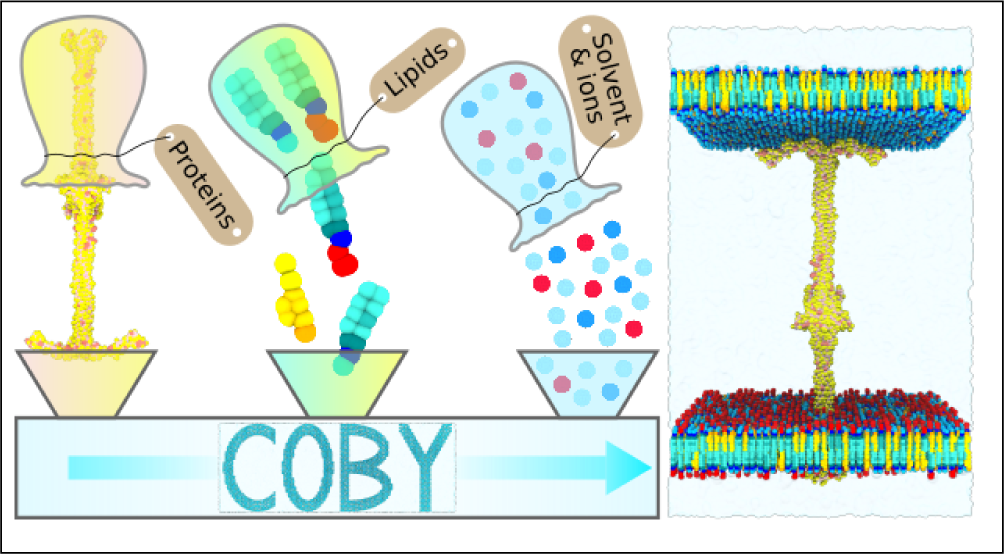

## 1 Introduction

Dynamics and function within a biological system are inextricably linked. Molecular dynamics (MD) simulations are a computational tool used to investigate motions within a macromolecular system on microsecond time-scales and in atom- or near atom-level resolution. By using experiment-informed structures paired with force fields defined by realistic inter-atomic potentials, simulations complement the experimental method and allow for a detailed view of an individual system in motion. ^1^

Continuously increasing computing resources, paired with the advancements in integrative biology, makes it possible to model systems of ever greater complexity. One particular area of interest is complex membranes, as membrane composition is linked to a plethora of biological functions - protein regulation through lipids, ^2–8^ modulated protein kinetics,^9^ protein localisation,^10^ and modulated biophysical properties of the membrane.^11^ However, computational costs of achieving adequate sampling increase dramatically with system complexity. Simulating the complex membranes in a coarse-grained (CG) resolution, where multiple interconnected atoms are grouped together to form a bead, allows for longer sampling of bigger systems, and ultimately a wider exploration of the conformational landscape ^12,13^ — a feat that is not easily achievable in atomistic simulations.

To run the MD simulations, one needs to start with system building, which is a user-involved process and can quickly become laborious in higher-complexity systems. The systems are therefore often built with the help of a software that automates the building process. It follows that system accuracy is directly dependent on the accuracy of the employed code. ^1^

Membrane building in particular is reliant on the available software.^13^ The procedure is non-trivial, as the membrane structure is not directly sourced from a given set of experimentally determined atom coordinates (as is the case with proteins); instead, the coordinates need to be built “from scratch”, based only on a few given parameters: membrane patch size, lipid types and their ratios, and the assigned area per lipid (APL). Ideally, all of the given parameters should exact to the given value in a built membrane; however, this is often not the case. Fundamentally, a membrane built for the purpose of MD simulations can only accept a whole number of lipids, which in consequence requires non-trivial consideration of the given parameters, as rounding can lead to value drift. This problem is exacerbated in complex membranes, where each lipid ratio (both within and between the leaflets) needs to be satisfied using a finite, integer number of lipids. If not considered, resulting ratios will be offset from the requested values and result in an inaccurate model of a complex membrane.

There exists a number of tools that can be used to automatise building of biomolecular systems (Charmm-GUI,^14–16^ TS2CG,^17^ Insane,^18^ packmol,^19^ polyply^20^), and while each offers a bespoke set of functionalities, we identified the need for an increased accuracy in terms of lipid ratios in complex membrane systems, coupled with ease of use, speed, and customisation ability.

Here we introduce a new system building software tool named COBY, which is an opensource Python package for building Martini coarse-grained systems. ^21^ COBY was primarily designed to build flat complex membranes while accurately handling the given lipid ratios, but it has been broadened to handle the creation of systems without a membrane component. It was designed to be fast and easy to use for a typical user, but also highly customisable in case of more specific system requirements. The software can be run both within a Python script or as a terminal command-line, and it uses a single command to specify all the system requirements.

It allows for easy creation of a broad spectrum of flat membrane-based systems: bilayers and monolayers of any given complexity, systems with any number of proteins (that can be of the same or different type), phase-separated membranes, stacked membranes, or membrane patches and pores of various shapes. COBY also handles building of the associated elements of the system, namely, membrane-inserted or soluble proteins, solvation (including partial solvation), and solute flooding. The package also includes developer-friendly features, such as easy handling of multiple parameter libraries, importing lipid structures to be used in the membrane building step, building molecules from fragments, and importing solute molecules for the purposes of flooding. Finally, while COBY defaults to “best practice” calculation methods in order to build the desired system, it also offers a set of alternative methods that can be used instead (i.e., inter-leaflet lipid optimisation options, multiple approaches to system neutralisation and salting, and handling of mixed solvent ratios).

The code, documentation, tutorials and license are available at github.com/MikkelDA/COBY.

## 2 Theory

The leading principles that guided the design of COBY were: i) accuracy in system building, particularly in terms of lipid ratios in complex membranes, system neutralisation, and solvent concentration; ii) versatility in code use, giving as high as possible level of freedom to the user to create systems of choice; iii) easy builds of special-case systems, which include stacked membranes, membrane patches and pores, or phase-separated membranes and solvents; and iv) advanced developer features, which feature molecule importers for the inclusion of new solute and lipid types, building of molecules from fragments, and a possibility to use multiple parameter libraries (with different versions of molecule types) that can be mixed and matched as desired.

This combination of versatile functions and new features ultimately allows for the user to build highly complex and biologically relevant systems in a single step. Importantly, careful consideration of accuracy in the building procedure assures that the produced systems are reliably built to the standard requested by the user.

### 2.1 Code syntax

The COBY package is written in Python 3 and the command can be used either within the Python script, or as a command line in the terminal window. The code is organised in a way where the entire simulation system is built in a single command call. Individual elements of the system are specified under bespoke arguments (e.g., membrane, solvent, protein) which by default accept a string argument. Within the string argument, sub-arguments can be used to precisely define and fine-tune individual system elements. Finally, there are environment arguments that deal with general COBY processing modes or output (e.g., backups, naming, output files, verbosity of the output, or setting up random seeds). An example dummy code in Python script syntax showcases the general structure of the command line:

`COBY.COBY(`

   `argument_1 = “SUBARG1:VAL1 SUBARG2:VAL2”,`

   `argument_2 = “SUBARG3:VAL3:SUBSUBARG4:VAL4”`

)

### 2.2 Code organisation

Arguments passed onto COBY are initially preprocessed, translating a string of sub-arguments into variables that can be used later by the program. Preprocessing ensures that the entire argument is read and understood before any core processing is done.

The preprocessed system elements are subsequently built in a serial fashion (Fig. 1), where individual elements of the system are added in a predefined order: first, the script creates a simulation box, then positions imported protein structures, followed by membrane creation, flooding with imported solute molecules, and ending with the solvation step.

**Figure 1:**
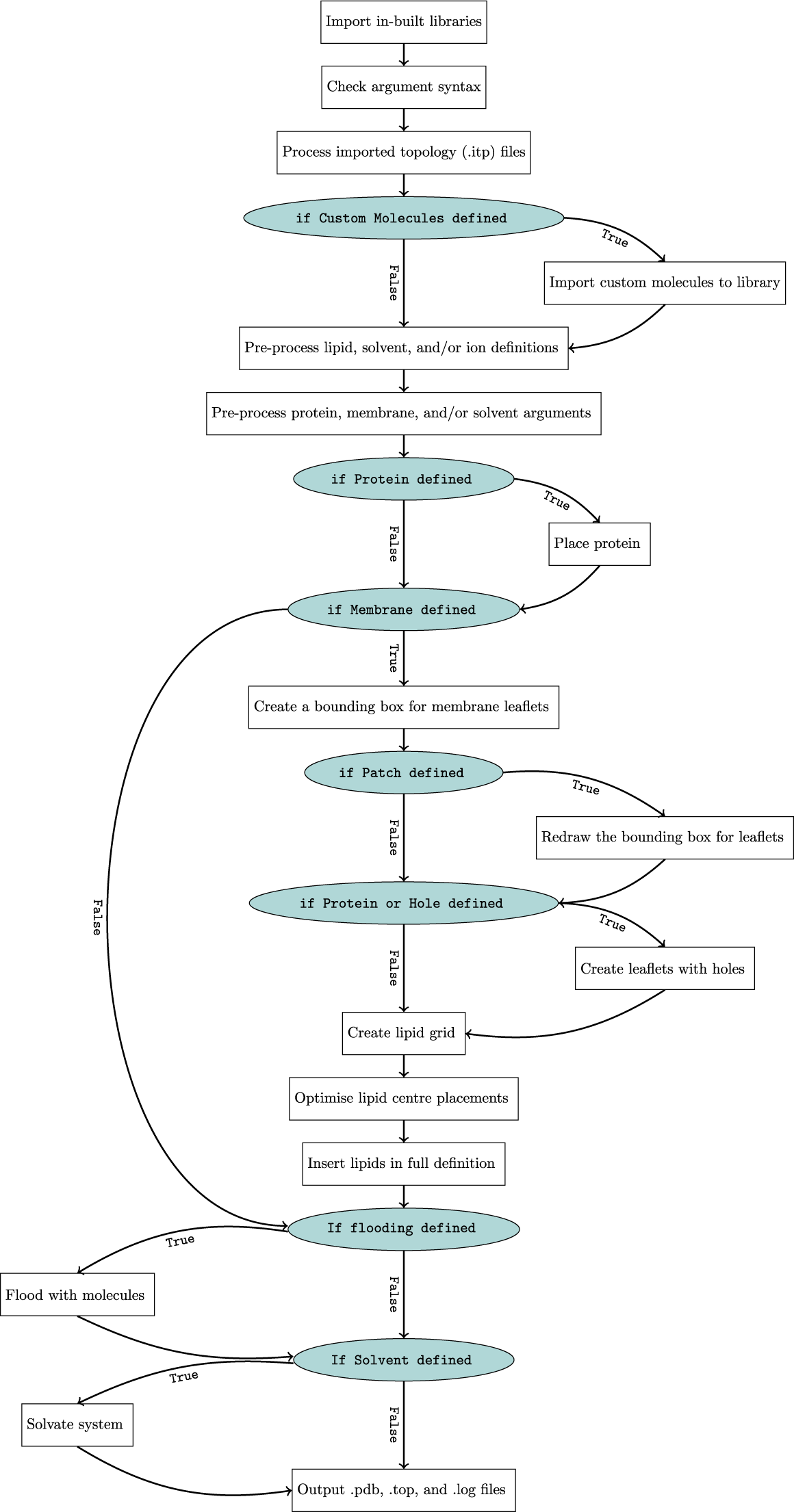
COBY workflow. The simplified code workflow shows the if-statements in teal circles, Boolean evaluations next to arrows, and the procedure steps in white boxes.

To run correctly, the code requires only the simulation box to be specified, alongside at least one structural element of the system (protein, membrane, solvent, or another imported structure of choice).

### 2.3 Membrane and protein treatment

COBY can create any number of membranes and insert any number of proteins. Both membranes and proteins can be placed at any given position within the simulation box, though membranes are restricted to the *xy*-plane, whereas proteins can be rotated and translated in any direction.

COBY treats leaflets as semi-independent components of the membrane. Processes relating to the membranes are calculated on a leaflet-by-leaflet basis, allowing for the creation of highly complex membranes and monolayer systems within the same argument-syntax framework.

When considering structures occupying the leaflet space (e.g., transmembrane proteins), leaflets are represented as rectangles with a height defined as the span of a tallest leaflet lipid and an additional buffer on each side. Otherwise, leaflets are represented as 2D rectangular polygons using the Python 3 module shapely.^22^

If a protein is placed in the proximity of a leaflet, then each protein bead is checked for potential overlap with the 3D leaflet space (Figure 2). By considering the bead overlap on a leaflet, rather than a membrane basis, the COBY lipid placement is better suited to accommodate irregularities in protein shape.

**Figure 2:**
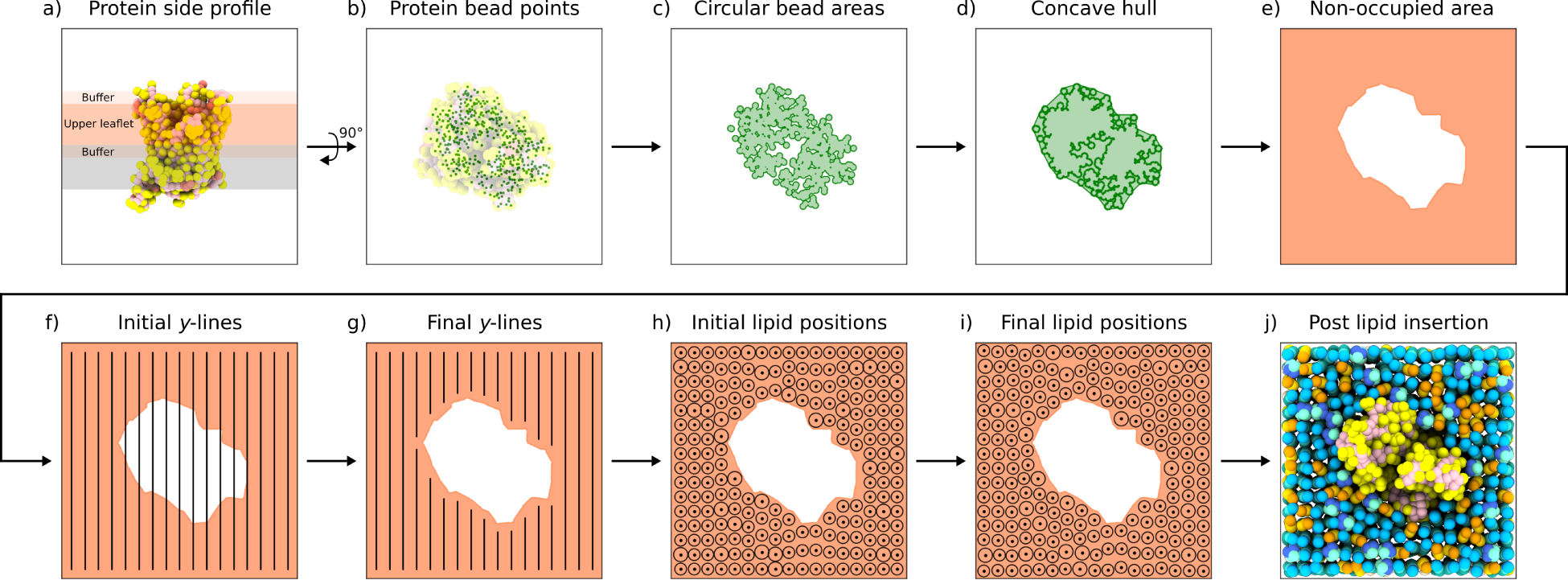
Membrane leaflet creation process in COBY. **(a)** Protein beads that are occupying leaflet and buffer spaces are **(b)** flattened onto a 2D surface as points. **(c)** The bead points are converted to circles using the Python module shapely and are merged if they are overlapping. **(d)** A concave hull is calculated from the surface points using the Python module alphashape, with an *α* value calculated from the diameter of the lipids present in the leaflet and an additional buffer distance. **(e)** A negative of the occupied space is used for lipid insertion. **(f)** A series of *y*-lines describing the possible placement of lipids is created. **(g)** The line segments that traverse the occupied areas of the leaflet are removed. The resulting line segments are then assigned a number of lipids that can be stringed along the *y*-line. If the total number of lipids in a leaflet is too small, then the density of the *y*-lines or the stringed lipids is increased, and the process is repeated. **(h)** All lipids are represented as circles with diameters corresponding to the specific lipid type. Lipid circles are inserted onto the grid in a random order, and **(i)** any overlaps are prevented by an internal optimisation algorithm. Finally, **(j)** the lipid and protein beads are placed in the 3D system.

COBY features tools for creation of membrane patches and artificial membrane pores. These can be made either by using predefined shapes, or by defining polygon points in the *xy*-dimension. pores are treated in the same manner as membrane-inserted proteins, and are thus considered as occupied areas in the membrane-building process. On the other hand, membrane patches are treated as membranes with a custom surface shape in the *xy* plane.

For large membranes, COBY segments the leaflets for faster processing. Similarly, if a membrane is segmented by the user or elements of the system (e.g., a protein spanning across the entire *x* or *y* direction, cutting the membrane in two), then the pr ogram will treat them as individual sub-leaflets contained within a leaflet, with the processes being divided accordingly.

### 2.4 Lipid packing

The lipid packing procedure involves determining an absolute number of lipids in leaflets, followed by an initial lipid placement, and finally the position optimisation. The allowed lipid occupancy area within a leaflet is calculated by taking the area of the whole leaflet and subtracting the area occupied by proteins or artificial pores. This area (*A*_free_) is then used to calculate the maximally allowed number of lipids (*N*_COBY,max_) from the area per lipid (APL) which is assigned by the user for each individual leaflet (or relies on a default value of 0.6 nm^2^). The result is then rounded to the nearest integer:

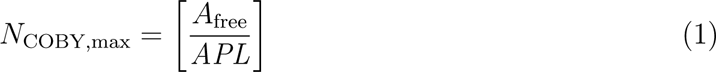

The *N*_COBY,max_ is then allocated across the requested lipid types, taking their internal ratios into account, with the number of each lipid type being rounded down to the nearest integer. A resulting sum of lipids is considered the minimal allowed number of lipids (*N*_COBY,min_) in the leaflet as

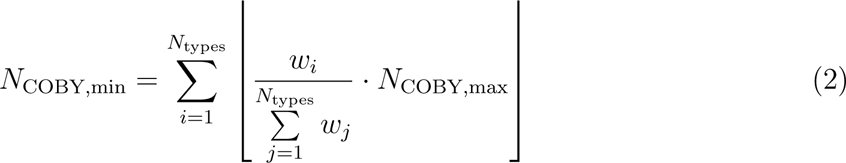

where *N*_types_ is the number of lipid types, *w* is the inter-lipid ratio of a given lipid, and *i* and *j* are the indices of the lipid types.

COBY includes multiple algorithms for optimising the absolute number of lipids in a leaflet. For instance, the default algorithm first adds *N*_COBY,min_ lipids which, as shown in Eq. 2, can be subdivided into a whole number of lipids of each type. Then, *N*_COBY,max_ is reached by iteratively adding lipids of the most underrepresented type relative to the ideal requested ratio.

Lipid placement is initiated by creating a grid within the non-occupied area of the leaflet, followed by optimising the line spacing informed by the final lipid number (Fig. 2e-g). Each lipid is represented in 2D as a circle, the diameter of which corresponds to the *xy*-plane diameter of that lipid type (in CG representation) and an additional buffer. Lipid circle centers are then positioned on the grid, and any overlaps between the circles are relieved by using a distance-based optimisation algorithm (Fig. 2h-i). In this step, the lipid circles are exerting a gentle pushing “force” to the neighbouring circles, causing them to rearrange. Overlapping circles are strongly pushed apart in order to prevent bead overlaps in the final system. A pushing “force” is also applied by the edges of the box and the membrane-occupied areas. With each step, the force is gradually reduced, and optimisation is completed when the maximum force on any lipid circle is smaller than a set tolerance value.

### 2.5 Treatment of solvation

Solvation steps in COBY are independent of the solvent molecule of choice. While the biological systems predominantly feature water as a solvent, the software can use any molecule as a solvent, as long as the structure and the molarity information are provided.

Solvation and flooding steps are performed in a 3D space, which is referred to as the solvation box. A solvation box is first rasterised by a 3D grid, with the grid resolution determined from the length of the largest solvent molecule. The grid points that overlap with other system beads (i.e., proteins and lipids) and the hydrophobic volume of a membrane are marked as unavailable, whereas the remainder can be used for solvent insertion.

An initial free volume of the box is calculated by taking the total box volume and subtracting the volume of all pre-existing beads (assumed to have a radius of a regular bead in Martini 3, *r* = 0.264 nm).^23^ This volume is then used to calculate a total number of solvent particles based on a given concentration. By contrast, flooding inserts an explicit number of particles in the system.

Solvent neutralisation is carried out in two steps, first by adding the desired concentration of ions, after which the ions are added or removed in order to neutralise the solvation box. COBY offers three neutralisation algorithms, where i) counterions are added, ii) co-ions are removed to reach neutralisation, and where iii) the addition of counterions and removal of co-ions is combined.

In mixed solvent cases, the inter-solvent ratios can be interpreted either as ratios based on the CG bead numbers, or as ratios that take into consideration the underlying atom-to-bead mapping (e.g., 4-to-1 for regular water beads or 3-to-1 for small water beads in Martini 3).

### 2.6 Stacked membranes

COBY features a specialised argument to create stacked membrane systems. It allows for the specification of an exact number of membranes and the distances between them. Intermembrane distances can be calculated either from the membrane centres, the maximal lipid heights, or the average lipid heights. Exact compositions can be set for each individual membrane and solvent space, allowing for an easy creation of complex systems.

### 2.7 Topology handling

COBY reads topology files and primarily uses them to obtain charge information for imported molecules. Topologies are unambiguously linked to the imported molecules by using the names specified in the [moleculetype] segment of the GROMACS topology file.

The #include statements within the topology file are recursively processed, allowing the user to supply a single file that contains all required topological information. The output topology file is updated with the specifications of the built systems and can directly be used in the simulation preprocessing steps in GROMACS.

### 2.8 Parameter libraries and molecule import

Molecular structures are organised in parameter libraries. Libraries can be attributed a type (i.e., lipid, positive ions, negative ions, or solvent libraries), and a name. Multiple libraries of the same type (but with a different name) can co-exist, and can be used to differentiate between multiple implementations of the same molecule. This feature is particularly interesting in the development context, where molecule parameters can be sourced from different development libraries. Mixing the parameters from different libraries within the same program call is allowed.

Within the libraries, parameters are structured in two interchangeable formats. The primary format developed for COBY is available for lipids, solvents, solutes and ions, where the specific bead names, their internal positioning and bead-specific charges are set for each molecule. This data format handles single- and multi-residue molecules.

The secondary data format, scaffolds, is inspired by the grid-system used in *insane*,^18^ and is available for lipids specifically. Here, generalised pseudo-structures containing bead-positions can be used to create a multitude of different lipids. Charges can be designated for each scaffold and for the specific lipids within the scaffold. The scaffolding data format is only available for single-residue lipids.

Imported molecules are stored using the primary data format. The charges for the imported molecules can be read either from the topology file (if provided), or need to be specified by the user. Note that imported lipids also require orientation information in order to be correctly placed in a membrane.

### 2.9 Building molecules from fragments

Inspired by *insane*’s functionality of building lipids from fragments, we designed COBY so it too can use internal fragment libraries to build the structures of molecules of interest (e.g., lipids with specific head, linker, and tail combinations). An existing set of fragments is provided as an in-built library, but the fragments can also be imported (as described in the previous section) and used in the building procedure. Notably, COBY can create lipids with complex profiles, as the molecules are dynamically built by joining the building blocks at the specific attachment points. For instance, the number of tails in COBY is dependent on the linker type and its number of attachment points, rather than the hard-coded number, which allows the user to import their own complex linkers and use them to create custom lipids with a number of tails that is not necessarily two. Note that COBY outputs structures, but not the topologies of the built molecules, and running the MD simulations will require the users themselves to provide the corresponding topology files to the simulation software.

molecule_builder can be used as an argument within COBY.COBY command line, where the built molecule can be directly passed onto the system builder and used in membrane, solvation, or flooding arguments. It is also possible to embed arguments for the molecule fragment builder in the topology files. The lines starting with; @COBY and followed by the molecule fragment builder arguments are read and interpreted by COBY, and the created molecules can then be used in a system building step.

Alternatively, molecules can be imported or created in a separate COBY.Crafter command call. COBY.Creator does not build the ready-to-simulate systems, but instead only outputs the structure of a requested molecule (either by obtaining it from an in-built library, or by building a molecule using fragments). By having an option to skip the system-building step, COBY can be used to straightforwardly provide structure files of the desired lipids, which is a routine requirement in the force field development process.

## 3 Methods

### 3.1 System showcase

Seven systems have been created to showcase the flexibility and versatility of COBY.

The code used to create the systems can be found in the Github repository under the Tutorial section (github.com/MikkelDA/COBY), and the system details are listed in Table S1.

The figures were visualised with ChimeraX version 1.8. ^24^

### 3.2 Code accuracy comparison

When reflecting on accuracy in membrane building, we focused on three considerations: the deviation of the total number of the inserted lipids from the ideal number of lipids; the deviation of the inter-leaflet lipid ratios in cases where the two leaflets have different APL values; and the deviation of the resulting lipid ratios from the requested ratios.

#### 3.2.1 Total number of lipids

The ideal number of lipids that should be inserted in a leaflet (*N*_ideal_) is calculated from the area of the leaflet and the APL.

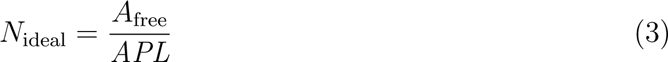

However, the constraints of the build — namely, that i) the system is built with an integer number of lipids ii) placed within a user-defined leaflet area and iii) with consideration of a given APL — necessitates a compromise between the given properties so that the resulting deviation from the relation of Eq. 3 is minimal.

We compared the accuracy of COBY and *insane* algorithms in reproducing the *N*_ideal_ by monitoring the deviation as a function of an increasing membrane size.

The number of lipids inserted by *insane* in a leaflet (*N_insane_*) relies on two rounding operations and is calculated by

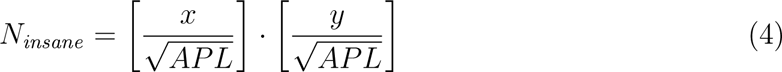

where *x* and *y* are the side lengths of a given membrane.

In contrast, the number of lipids inserted by COBY (*N*_COBY,max_) is shown in Eq. 1.

A comparison of software accuracy based on the deviation from the requested lipid ratios was based on concurrent command calls to COBY and *insane* with an increasing box size (between 4 and 51 nm and by 0.05 nm increments) and with both leaflets having APLs of 0.6 nm^2^.

#### 3.2.2 Inter-leaflet lipid ratios

The deviations from the *N*_ideal_ within a leaflet also affect the accuracy of the inter-leaflet lipid ratios if the two leaflets have differing APL values. We assessed a deviation of the inter-leaflet lipid ratios between the two software tools as a function of the membrane size, where the upper leaflet was assigned an APL of 0.6 nm^2^, and the lower leaflet an APL of 0.45 nm^2^.

#### 3.2.3 Intra-leaflet lipid ratios

In a complex membrane, different lipid types are assigned to a given leaflet using ratio specifications. Considering that the number of lipids always needs to be an integer, the assignment of lipids across types becomes a non-trivial issue. The building algorithms need to minimise the errors by considering a number of requirements that, taken together, often do not have an analytical solution. The COBY algorithm that optimises the lipid assignment across types is presented in the Theory section. The insane algorithm, on the other hand, assigns the number of lipids across types by using a rounding down operation.

### 3.3 Speed tests

The speed of COBY and *insane* has been assessed by calculating the mean time to create a system over 5 attempts. Three series of systems have been created. The first contains only a membrane, the second contains only solvent, and the third includes both a membrane and solvent. The membranes are assigned only POPC lipids with an APL of 0.6 nm^2^. The solvent contains water and 0.15 M NaCl. The *z* dimension of a box was kept constant at 10 nm, while the *x* and *y* box size lengths were serially increased from 4 to 51 nm by 0.05 nm increments.

## 4 Results

We developed COBY, a versatile Python-based software tool for creating Martini CG systems that can be pipelined into MD simulations. The software includes a range of functionalities for easy creation of any number of flat complex membranes, including membrane patches, pores, monolayers, and stacked membranes. Additionally, COBY handles solvation, ion concentration and neutralisation, allowing for multiple solvent compartments of different compositions. The structure and topology importers can be used to introduce molecules not present in the in-built libraries, e.g., proteins, ions, ligands, solutes or lipids. For simplicity, single structures are imported as “proteins” in the command line, but in practice any molecular structure can be imported using the same syntax.

### 4.1 Versatility of the built systems

The increase in system complexity has been a trend in the modelling community as a way to better reflect the native biological environment. COBY allows for fast and effective creation of such demanding systems, which had previously been handled by non-standardised procedures based on manual “system-stitching”. We structured COBY in a way that all the elements of the system can be specified in a single command call, meaning that the program can optimise the building procedure with regards to the requested components. This, in effect, can produce a great variety of high-complexity systems in a single step (Fig. 3).

**Figure 3:**
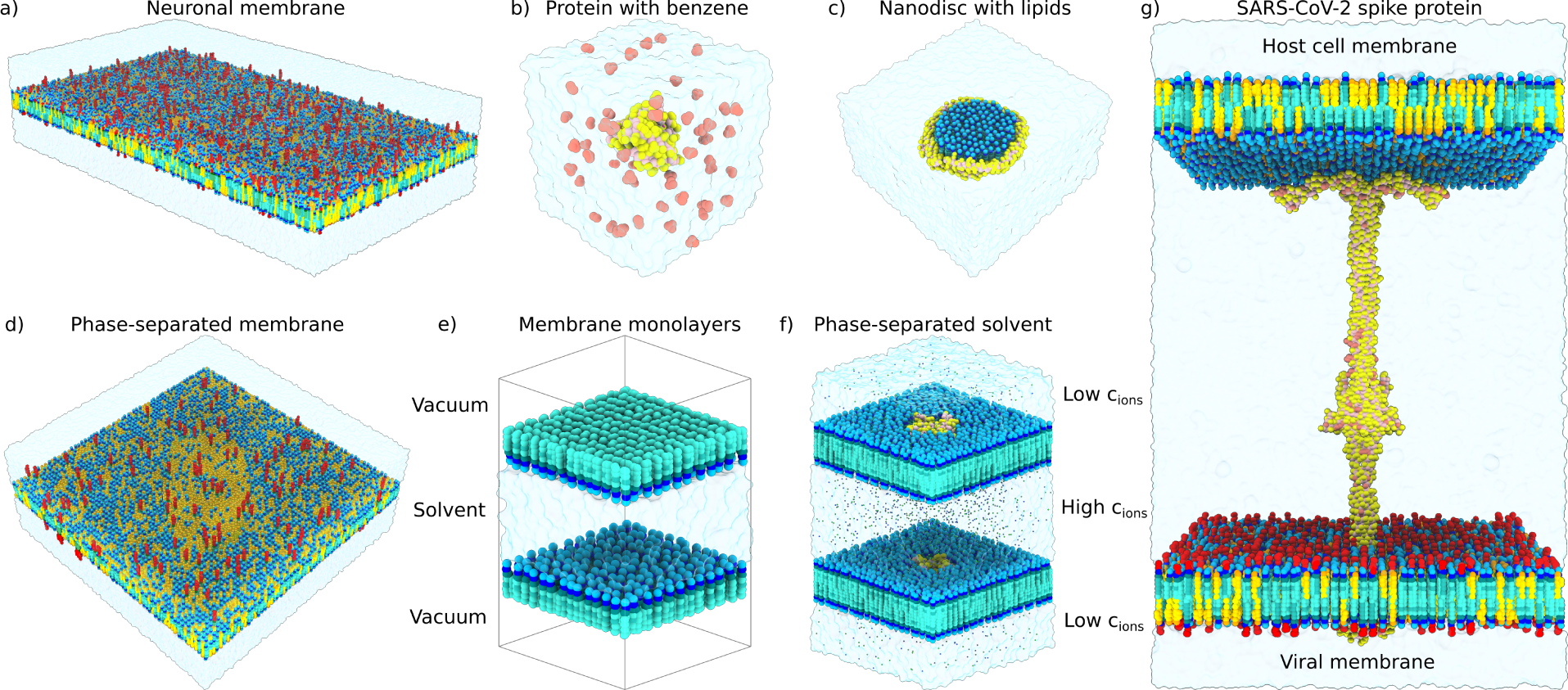
Showcase of systems built with COBY and using a single commandline call. **(a)** Complex asymmetric neuronal membrane consisting of 58 different lipid types described by Martini 2 mapping. Based on Ingólfsson et al. ^25^ **(b)** A benzene flooding setup, with a random placement of benzene in the solvent surrounding a human K-Ras protein (PDB: 4OBE).^27^ **(c)** A nanodisc in solvent, consisting of two apolipoproteins (PDB: 6CLZ)^28^ enclosing a DMPC (1,2-dimyristoylphosphatidylcholine) membrane patch. **(d)** A phase-separated giant unilamellar vesicle membrane. An ellipsoid liquid-ordered phase patch is surrounded by a liquid-disordered membrane. Lipid compositions are based on Hammond et al.^26^ **(e)** A monolayer simulation setup, consisting of two monolayers in a mirrored orientation, a solvent bridging the headgroup space, and vacuum across the tail region. **(f)** Voltage-gated potassium channels (PDB: 3F5W) ^29^ with a modelled transmembrane potential; two stacked membranes separate two solvent regions with a high (0.4 M) and physiological (0.15 M) NaCl concentration in the central and edge solvent space, respectively. The system is inspired by Kutzner et al. ^30^ **(g)** An extended intermediate of a SARS-CoV-2 spike protein spanning the viral and host membrane. The system was modelled using the spike conformation from Su et al. ^31^ The host membrane was based on the simplified average membrane,^25^ and the viral membrane composition was informed by the available SARS-CoV-2 lipidomics data.^32^

One example of a demanding system often built in CG resolution are complex membranes (Fig. 3a). In our example, we recreated a neuronal membrane model from Ingólfsson et al. ^25^ that contained 58 different lipid types. On top of being highly complex in terms of lipid species, the neuronal membrane was also asymmetric, which was reflected in different leaflet APL values. COBY handled the lipid type assignments in correct ratios while taking into consideration the leaflet APL values. All lipid types were imported from a Martini 2 lipid library file. The entire membrane building procedure and solvation took under 30 seconds on an office workstation.

COBY can also perform flooding of systems with user-imported solutes, exemplified by the benzene flooding setup of the K-Ras protein, relevant for uncovering cryptic pockets (Fig. 3b). Both K-Ras and benzene were imported from their respective structure files, and benzene was added to random positions and orientations to the solvent in user-specified concentration, while ensuring no overlaps with the protein. COBY correctly handled solvent and ion concentrations even after the addition of solute molecules, as it considers only the unoccupied volume for calculating the required number of solvent and ion particles.

Nanodics are a synthetic membrane constructs that can be used for the *in vitro* studies of membrane proteins. COBY was used to create a nanodisc in solvent (Fig. 3c), where a specialised membrane subargument was used to specify lipid placement within the borders of the apolipoprotein, as opposed to the default lipid placement surrounding the protein. The code also handled the solvation of the remainder of the solvation box unoccupied by the nanodisc.

Furthermore, COBY can be used for creating phase-separated membranes, as exemplified by the model of a giant unilamellar vesicle membrane (Fig. 3d). This system was inspired by the experimental setup by Hammond et al. ^26^ that featured liquid-liquid phase separation. This system was modelled by combining the hole and patch subarguments that create regions of distinct membrane compositions of any given shape (due to its in-built polygon and curved shape functionality).

On top of making the lipid bilayers, COBY can easily handle monolayers, as the code by default performs virtually all membrane building steps in the leaflet subspace. The monolayer setup used for MD simulations features two monolayers facing each other, a layer of solvent between them, and the vacuum region spanning the tails (Fig. 3e). As COBY can handle any number of membranes (leaflets) and solvent spaces, the system was built in a single step by specifying the offset between the two monolayers and the solvent composition.

Multilayered membrane systems can also be created more stringently by using a special stacked_membranes argument. Stacked membrane systems are often used for simulating transmembrane potential conditions, as they feature multiple isolated solvent spaces and solvents of different compositions. In this example, two voltage-gated ion channels — one in each membrane — were positioned across two solvent spaces, with the extracellular segments facing the high-ion concentration solvent, and the intracellular components towards the low-ion concentration solvent (Fig. 3f). One of the proteins had to be rotated 180° the *x*-axis, which was also performed by COBY, as it handles protein rotation, translation, and insertion into the membranes (Fig. 2), and does it for any number of proteins.

The composition of individual membranes in a stacked membrane system can also be explicitly defined. Furthermore, the placement of proteins is not restricted to the membrane subspace — instead, they can either be fully soluble (as showcased in Fig. 3b) or they can span multiple membranes. We built a system featuring an extended spike protein of SARSCoV-2 spanning the viral and the host cell membrane (Fig. 3g). This extended intermediate is formed when the viral fusion peptides are inserted into the host cell membrane and preclude the viral-host membrane fusion event essential for successful viral entry into the cell. COBY built this system in a single command call by creating two complex membranes with different compositions, and by positioning the extended intermediate so that it spans the solvent space between the two membranes. The system creation took around 10 seconds on an office workstation.

### 4.2 Other algorithm features

During code development, a particular attention was paid to the customisability of the building procedure, which also includes a choice of algorithms relating to system neutralisation, mixed solvent specification, lipid ratio handling, and lipid packing procedures. Additionally, we implemented several developer-friendly features. Namely, these include multiple parameter libraries that can be mixed-and-matched where necessary, and a lipid builder that can create lipid structures from an input from fragment or topology files, allowing the user to create their own lipids of choice.

The neutralisation procedures allow the user to exert more control on the types of ions used to neutralise the system, which can be of great value if the exact concentrations of all ionic species in the system are of interest in the study. While formulating the mixed solvent ratios, the user has the freedom to choose between several different options of interpreting the given ratios — namely, if the ratios are considered in a CG or atomistic resolution. Similarly, calculating the absolute number of lipids based on lipid ratios requires trade-offs in accuracy, bearing in mind that the final number of lipids always needs to be an integer. Prioritising the accuracy of APL as opposed to the lipid ratios, or vice-versa, can be controlled by the user. Finally, the fine-tuning of lipid placement is performed by an optimisation algorithm. It is employed in cases of overlap between the lipids after the initial placement. However, the users can opt to apply the optimisation step in all membrane builds, regardless of the overlap criterion.

Molecule parameterisation procedures require ample testing of the designed parameter sets, which might also involve alternative molecule mapping schemes. With that in mind, we implemented a library system of molecule types, where molecule structures (and charges) are saved within a particular library. The libraries can be mixed and matched during the build procedure, allowing the user full flexibility in their choice of system molecules. Additionally, the users can easily import new molecules (i.e., solutes, lipids) from structures with a molecule_import argument and use them as a part of the building procedure (e.g., imported lipids can be included as membrane components or imported solutes as solvent components). Lipids can also be manually constructed using a molecule_builder argument from a set of library fragments, either by specifiying it within the system-building COBY.COBY command, molecule-building COBY.Crafter command, or within a topology file.

### 4.3 Accuracy of membrane builds

As previously outlined, the constraints of the membrane building procedure — the need to accommodate APL, lipid ratios, and a membrane patch size with an integer number of lipids — inherently lead to an error within one or more of the requested values describing the membrane properties. COBY was designed with the aim of minimising the build errors with a protocol described in the Theory section.

In order to test the accuracy of membrane building, we examined the effect of membrane patch size on the deviation from the ideal number of inserted lipids, as dictated by leaflet APL. We compared the performance of COBY with a commonly used CG membrane building software, *insane*. The deviation from the ideal number of lipids was appreciably higher for the membranes built by *insane* than it was for the membranes generated by COBY (Fig. S1a). The deviation for *insane*membranes was especially large in small lipid patches and it tangentially reduced with the membrane size, but remained observable even in the patches that are larger than 50 nm *·* 50 nm. On the other hand, the deviation from the ideal lipid number in the membranes built by COBY dropped below 1% at membrane patch sizes of 6 nm *·* 6 nm and remained under that value for all larger membrane patches tested.

We also assessed how the two algorithms compare in terms of preserving lipid ratios between the two leaflets in cases where the requested leaflet APLs are not the same (Fig. 4b). Similarly to the previous example, COBY rapidly minimised the errors and remained very close to the ideal ratio, with only slight deviations visible for the smaller membrane patches. *insane* was comparatively producing much bigger deviations from the requested ratios that lessened with the increasing membrane patch size, but remained a noticeable factor affecting accuracy even in very big membranes.

**Figure 4:**
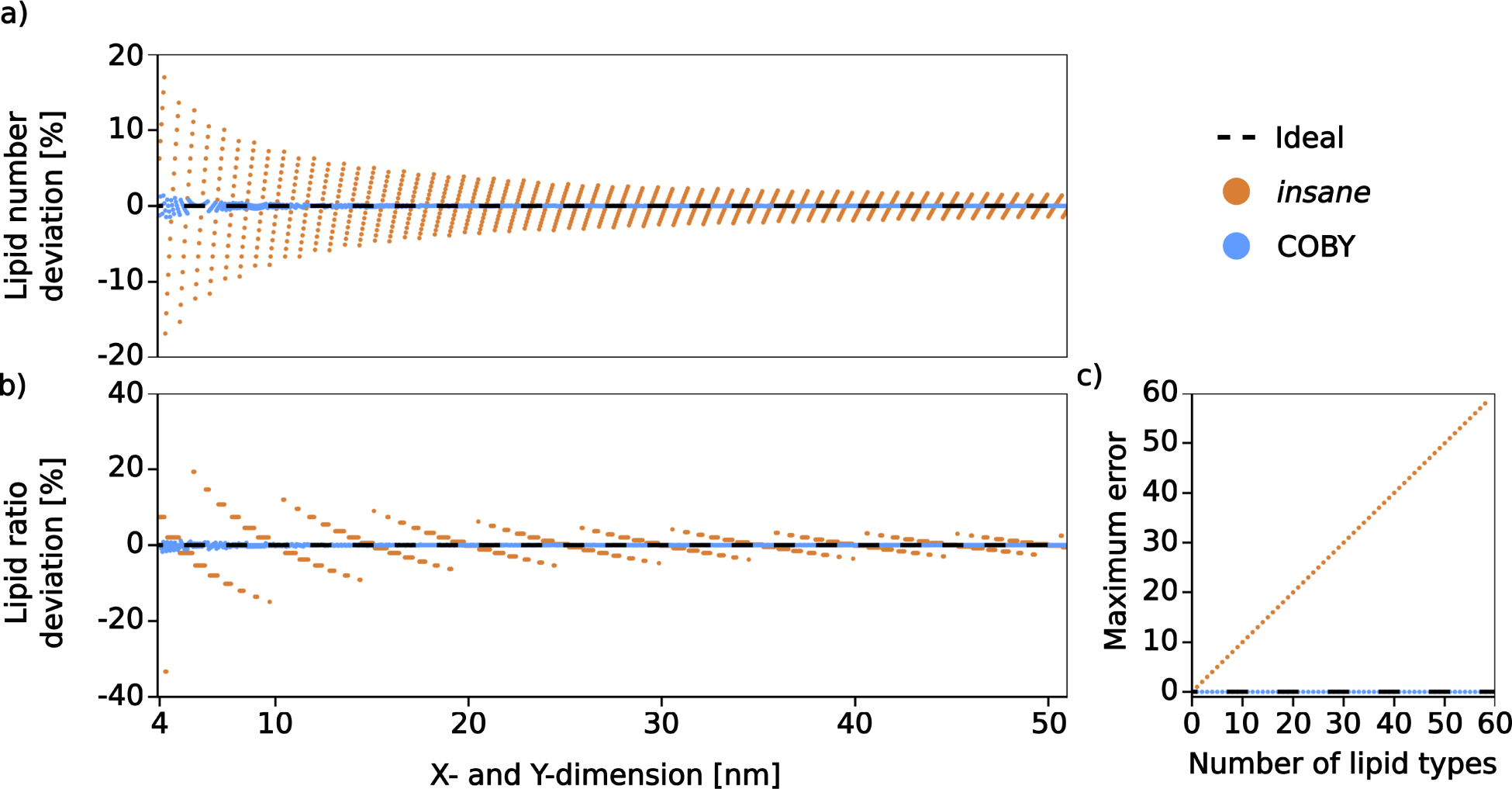
Comparison of accuracy in the membrane building step. The deviation from the ideal value (shown as a black dashed line) is shown for COBY (in blue) and *insane* (in orange). **(a)** The deviation in the number of lipids within a built leaflet compared to an idealised leaflet, expressed as a % deviation, and calculated as a function of an increasing membrane size (APL = 0.6 nm^2^). **(b)** The deviation in the ratio of lipid numbers between the two leaflets with different APL values (APL_lower_ = 0.6 nm^2^; APL_upper_ = 0.45 nm^2^) as a function of an increasing membrane size. **(c)** The largest possible error that can be accrued with the increasing number of lipid types within a leaflet.

These differences in accuracy are due to the architecture of the underlying algorithms. COBYs algorithm attempts to bring the number of lipids as close to *N*_ideal_ as possible, and the variations are mainly confined within the decimal values. As the number of inserted lipids is by necessity an integer number, these deviations are only observable in very small membrane patches and quickly disappear with an increasing membrane size. *insane* is comparatively more sensitive to the membrane size effects, as it assigns the number of lipids within a strictly defined 2D-grid (where the number of rows *n*_rows_ and the number of columns *n*_columns_ are informed by the *x*- and *y*-axis patch dimensions, respectively). With an increasing patch size, instead of expanding the membrane with just the required number of missing lipids, *insane* adds the entire rows and columns of lipids, which results in large deviations and jumps away from the ideal. The lessening of this effect observed with an increasing patch size can be attributed to the fact the the total number of lipids in *insane* scales with *n*_columns_ *· n*_rows_, while the error in the number of lipids scales with *n*_columns_ + *n*_rows_.

When building complex membranes, *insane* can accumulate errors as a result of improper handling of intra-leaflet lipid ratios (Fig. 4c). After calculating the required number of lipids of each type within a leaflet (dictated by user-requested ratios), *insane* software achieves an integer number by always rounding down, which can lead to a skewed ratio that worsens with each additional lipid type. This artefact is not present in COBY.

One aspect where COBY is inferior to *insane* is speed (Fig. S1). COBY takes more time to create the same type of systems due to the use of more complex algorithms that handle the system building procedure. In particular, solvent insertion is COBY is slower than in *insane*, primarily because COBY takes into consideration free volume and solvent molarity in order to calculate the required number of solvent molecules. In comparison, solvation in *insane* is handled by creating a simple 3D grid with spacing that roughly corresponds to the density of regular water beads. The water beads are then placed within the cells if there is no overlap with other elements of the system. The advantage of COBY’s approach is that solvation is not restricted to water — instead, any solvent molecule (that either exists in the library or is imported by the user) can be used to solvate the system, provided solvent-specific molarity. The apparent drawback of this approach, however, is speed. We decided to maintain the approach where we employ slower, more complex algorithms in order to create systems of choice, as they result in the systems that more accurately reflect the conditions requested by the user (Fig. 4). In other words, the trade-off between accuracy and versatility vs. speed was deemed a necessity, particularly when considering that even under these constrains, the creation of the larger systems (Fig. 3a,g) took under 30 seconds on an office workstation.

## 5 Discussion

We created COBY, an open-source software for building CG systems for MD simulations, with a primary focus on the accuracy of the building procedure, followed by the versatility of the produced output. The code development was primarily motivated by the need for accurate algorithms for system building. Under this premise, COBY was designed to minimise errors that stem from the inherent trade-off required to satisfy the user-requested system properties with an integer number of molecules. This is of particular importance in the membrane building procedures. For now, the code has been designed to create systems with Martini CG models. However, the underlying code infrastructure is designed so it can potentially be expanded to include other CG models (e.g., SIRAH^33^).

COBY demonstrates a greatly improved accuracy in system building, employing error minimisation algorithms which results in systems that accommodate the properties requested by the user. This, as we highlighted in the Results, is not a trivial task. If not handled with abject accuracy in mind, the system building software can generate errors that are difficult to predict, as the deviations from the ideal are neither well-documented nor linear. The result is a system that does not exactly correspond to the user request, and depending on the required level of precision and the exact system details, the significance of these errors will vary drastically on a case-to-case basis. This is also expected to affect the replicability of simulation studies, as unassuming changes in the starting conditions (e.g., simulation box size) can lead to unexpected errors in the built systems (incorrect APL values of the membrane). The improvements implemented in COBY allow for an accurate creation of multi-component systems, e.g., complex membrane models. ^25,34,35^ This alleviates the burden on the user to check, detect, and correct the errors generated by the software.

We designed COBY with complex systems in mind, and as such the software is able to handle a great variety of user requests within a single command line. We deemed this to be of particular importance, as the trends in the field of MD simulations are geared towards creating systems of ever increasing size and complexity. This is crucial in high-throughput protocols which utilise MD simulations, as they rely on the automated procedures to execute the entire pipeline,^36^ and should ideally not be restricted by the limited complexity of the available system building methods. Furthermore, programmatically-built complex systems hold value not only because they eliminate the need to perform a laborious task of “system-stitching”, but also because the procedure provides a straightforward approach to reliably replicate the simulation experiment. We concede, though, that COBY is designed to only handle flat membranes, and the curved membrane systems can instead be created with the TS2CG software;^17^ similarly, the generation of disordered polymeric assemblies (in the absence of a membrane) could be better handled with polyply, ^20^ as it is designed to generate the polymers in different starting conformations, while COBY treats the structure elements as rigid system components.

We structured the code syntax in a way that is easy to use and understand, while keeping up with the “spirit” of Python code structure familiar to most members of the simulation community. In particular, we reduced the number of compulsory parameters to a minimum, allowing for the quick and easy creation of simple systems. At the same time, we included a high degree of flexibility in terms of parameter selection, allowing the user to fine-tune the command according to their specific system requirements. We considered this of importance as the complexity of systems can arise in many different ways, and allowing the user to change a plethora of under-the-hood variables can help generate a desired setup, if the user is familiar enough with the software. Possible implementation of COBY within a graphical user interface environment, such as MAD,^37^ could further simplify the system-building procedure for many users.

Force field development is reliant upon extensive testing of the new parameters. Streamlined testing protocols depend on the availability of multiple versions of the same molecule under different mapping schemes. COBY parameter libraries are organised in a way that makes it possible to use multiple libraries within the same command call, allowing the user to mix and match the parameters of choice. Additionally, the user can create new libraries either by directly importing them, or by importing individual molecules from structure files and optional topology files. The information stored in the topology files is crucial for correctly treating the system charges. We therefore paid special attention to the implementation of a topology reader within COBY, as programmatic neutralisation schemes are crucial for streamlined system building. In addition to molecule importing functionality, a big portion of lipids can be built in COBY from fragments available from the in-built libraries. All of these functionalities, albeit not necessarily vital for the regular user, are nonetheless an important feature for the force field development process.

COBY is a tool that is meant to have a broad user base, as the simple systems can be built with a very few mandatory arguments — namely, box determining the size of the system, and at least one other element (e.g., membrane, solvent, or protein), while at the same time, it offers a great deal of flexibility to the experienced user wishing to employ it for complex system building or systematic parametrisation procedures. The code, several tutorials, a cheat sheet, and a detailed documentation is hosted on the project page at github.com/MikkelDA/COBY.

## Author Contributions

MDA: Conceptualisation, code writing and maintenance, manuscript — original draft & editing, user guides — original draft & editing;

PCTS: Supervision support, manuscript - review & editing;

BS: Funding acquisition;

LZ: Conceptualisation, supervision, project administration, manuscript — original draft & editing, user guides — original draft & editing.

## Acknowledgement

We thank Lisbeth R. Kjølbye for helpful discussions regarding desired software functionalities, Kasper B. Pedersen for exciting discussions, encouragement, and useful advice concerning lipid treament, Luís Borges-Araújo and Fabian Grünewald for astute discussion points in terms of desired developer features and software naming, Guillaume Launay for helpful advice towards improving code organisation, and Helgi I. Ingólfsson for providing the original lipid dataset for creating a complex neuronal membrane. P.C.T.S would like to thank the support of the French National Center for Scientific Research (CNRS) and the funding from research collaboration agreements with PharmCADD and Sanofi. P.C.T.S. also acknowledge the support of the Centre Blaise Pascal’s IT test platform at ENS de Lyon (Lyon, France) for the computer facilities. The platform operates the SIDUS solution developed by Emmanuel Quemener.^38^

## Supporting Information Available

Supporting information available: COBY workflow; a table detailing the showcase systems (Fig. 3); and the speed test comparison.

## Supplementary Information

**Figure S1:**
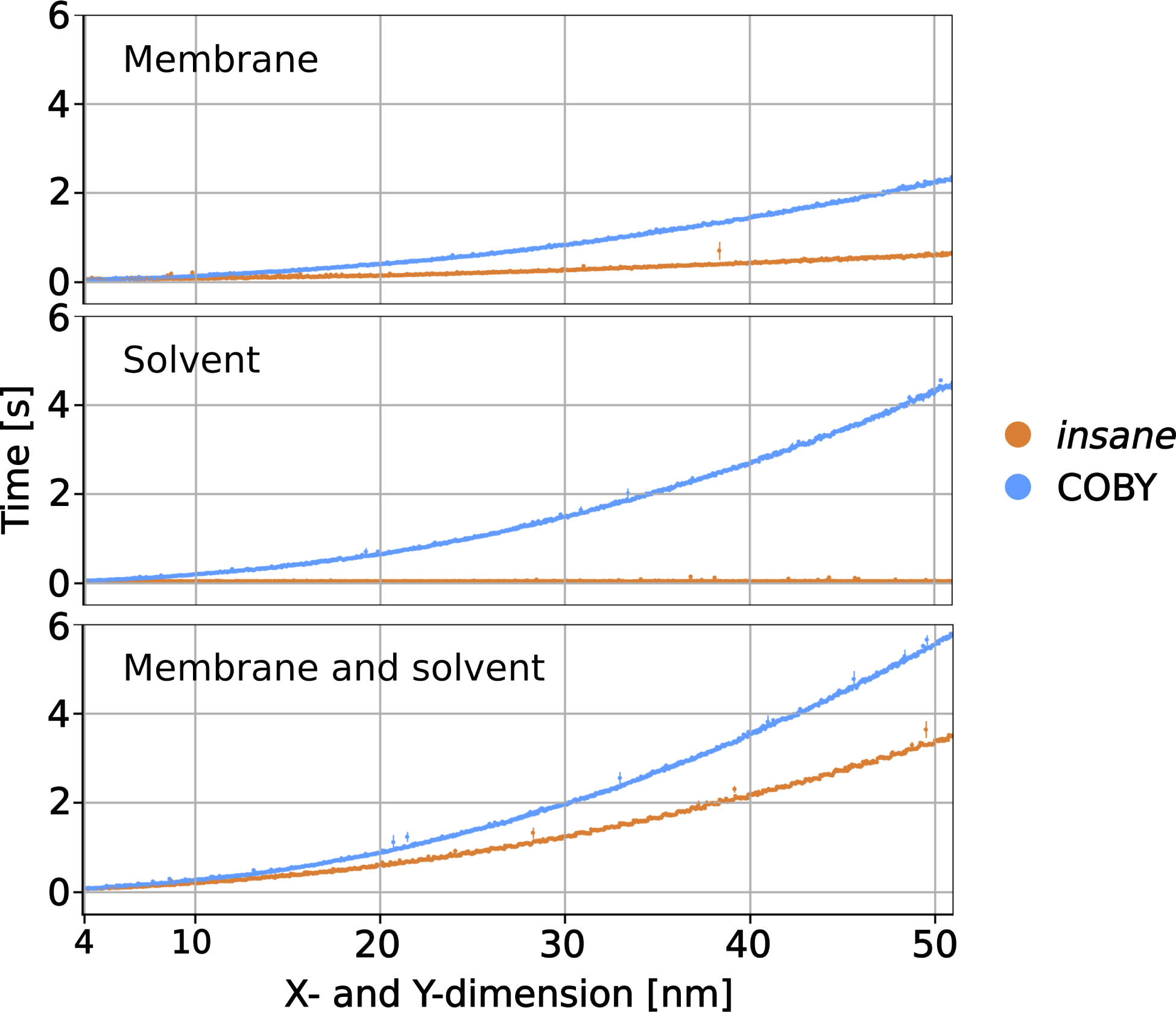
Speed comparison between COBY and *insane*. COBY performance is shown in blue, and *insane* in orange. The speed tests were performed on systems of an increasing size in the *x* and *y* direction, while the *z*-dimension was kept constant at 10 nm. Each data point was tested five times and is shown as the average and the standard error. The tests were performed on systems with only the membrane, only the solvent, and the combination of both. The membranes consisted of POPC lipids (APL = 0.6 nm^2^), while the solvent contained water and 0.15 M NaCl. Following the best-use practice, COBY was run from a Jupyter notebook, and the speed tests did not include the time required to import packages (which takes additional 3-4 s, depending on the available hardware). *insane* was run from a terminal.

**Table S1:**
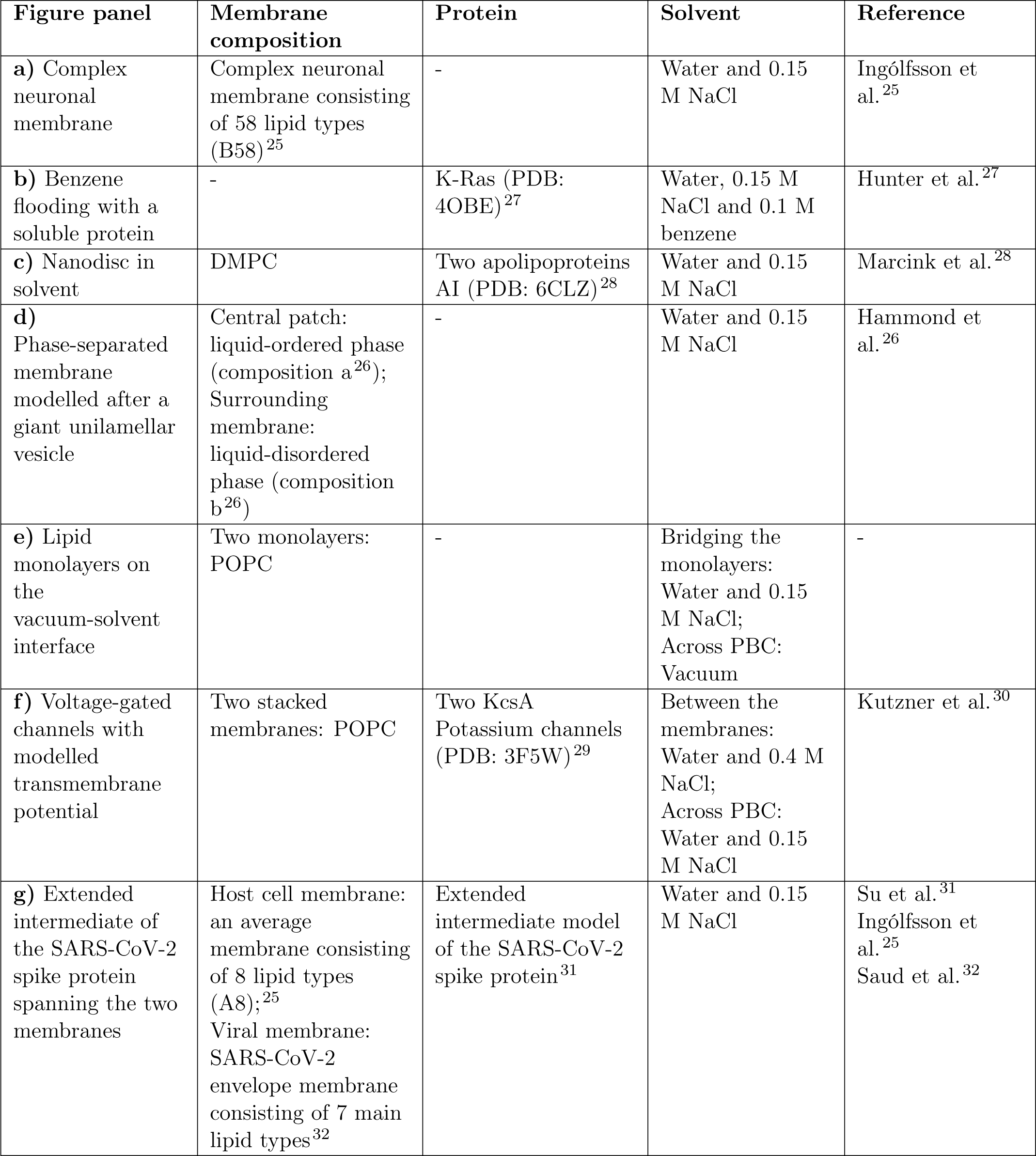
Overview of the built systems showcased in Fig. 3 with details of membrane, protein, and solvent composition. 1,2-dimyristoylphosphatidylcholine (DMPC); 1-Palmitoyl-2-oleoylphosphatidylcholine (POPC).

## References

(1) Hollingsworth, S. A.; Dror, R. O. Molecular Dynamics Simulation for All. Neuron 2018, 99, 1129–1143.

(2) Kervin, T. A.; Overduin, M. Regulation of the Phosphoinositide Code by Phosphorylation of Membrane Readers. Cells 2021, 10, PMID: 34069055.

(3) Niggli, V. Regulation of protein activities by phosphoinositide phosphates. Annu. Rev. Cell Dev. Biol. 2005, 21, 57–79, PMID: 16212487.

(4) Borges-Araújo, L.; Souza, P. C. T.; Fernandes, F.; Melo, M. N. Improved Parameterization of Phosphatidylinositide Lipid Headgroups for the Martini 3 Coarse-Grain Force Field. Journal of Chemical Theory and Computation 2022, 18, 357–373.

(5) Medoh, U.; Abu-Remaileh, M. The Bis(monoacylglycero)-phosphate Hypothesis: From Lysosomal Function to Therapeutic Avenues. Annual review of biochemistry 2024,

(6) Duncan, A. L.; Corey, R. A.; Sansom, M. S. Defining how multiple lipid species interact with inward rectifier potassium (Kir2) channels. Proceedings of the National Academy of Sciences of the United States of America 2020, 117.

(7) Kalinichenko, L. S.; Kornhuber, J.; Sinning, S.; Haase, J.; Müller, C. P. Serotonin Signaling through Lipid Membranes. 2024.

(8) Drew, D.; Boudker, O. Ion and lipid orchestration of secondary active transport. Nature 2024, 626, 963–974.

(9) Nilsson, T.; Lundin, C. R.; Nordlund, G.; Ädelroth, P.; von Ballmoos, C.; Brzezinski, P. Lipid-mediated protein-protein interactions modulate respiration-driven ATP synthesis. Sci. Rep. 2016, 6, 24113, PMID: 27063297.

(10) Jiang, Y.; Thienpont, B.; Sapuru, V.; Hite, R. K.; Dittman, J. S.; Sturgis, J. N.; Scheuring, S. Membrane-mediated protein interactions drive membrane protein organization. Nature Communications 2022, 13, 7373, PMID: 36450733.

(11) Levental, I.; Levental, K. R.; Heberle, F. A. Lipid Rafts: Controversies Resolved, Mysteries Remain. Trends in Cell Biology 2020, 30, 341–353.

(12) Corradi, V.; Sejdiu, B. I.; Mesa-Galloso, H.; Abdizadeh, H.; Noskov, S. Y.; Marrink, S. J.; Tieleman, D. P. Emerging Diversity in Lipid–Protein Interactions. Chemical Reviews 2019, 119, 5775–5848, PMID: 30758191.

(13) Marrink, S. J.; Corradi, V.; Souza, P. C. T.; Ingólfsson, H. I.; Tieleman, D. P.; Sansom, M. S. P. Computational modeling of realistic cell membranes. Chem. Rev. 2019, 119, 6184–6226, PMID: 30623647.

(14) Jo, S.; Kim, T.; Iyer, V. G.; Im, W. CHARMM-GUI: A web-based graphical user interface for CHARMM. Journal of Computational Chemistry 2008, 29, 1859–1865.

(15) Qi, Y.; Ingólfsson, H. I.; Cheng, X.; Lee, J.; Marrink, S. J.; Im, W. CHARMM-GUI Martini Maker for Coarse-Grained Simulations with the Martini Force Field. Journal of Chemical Theory and Computation 2015, 11, 4486–4494.

(16) Hsu, P.; Bruininks, B. M. H.; Jefferies, D.; de Souza, P. C. T.; Lee, J.; Patel, D. S.; Marrink, S. J.; Qi, Y.; Khalid, S.; Im, W. CHARMM-GUI Martini Maker for modeling and simulation of complex bacterial membranes with lipopolysaccharides. Journal of Computational Chemistry 2017, 38, 2354–2363.

(17) Pezeshkian, W.; König, M.; Wassenaar, T. A.; Marrink, S. J. Backmapping triangulated surfaces to coarse-grained membrane models. Nat. Commun. 2020, 11, 2296.

(18) Wassenaar, T. A.; Ingólfsson, H. I.; Böckmann, R. A.; Tieleman, D. P.; Marrink, S. J. Computational Lipidomics with insane: A Versatile Tool for Generating Custom Mem-branes for Molecular Simulations. Journal of Chemical Theory and Computation 2015, 11, 2144–2155, PMID: 26574417.

(19) Martínez, L.; Andrade, R.; Birgin, E. G.; Martínez, J. M. PACKMOL: A package for building initial configurations for molecular dynamics simulations. Journal of Computational Chemistry 2009, 30, 2157–2164.

(20) Grünewald, F.; Alessandri, R.; Kroon, P. C.; Monticelli, L.; Souza, P. C. T.; Marrink, S. J. Polyply; a python suite for facilitating simulations of macromolecules and nanomaterials. Nature Communications 2022, 13, 68.

(21) Marrink, S. J.; Monticelli, L.; Melo, M. N.; Alessandri, R.; Tieleman, D. P.; Souza, P. C. T. Two decades of Martini: Better beads, broader scope. WIREs Computational Molecular Science 2023, 13.

(22) Gillies, S.; van der Wel, C.; Van den Bossche, J.; Taves, M. W.; Arnott, J.; Ward, B. C.; others Shapely. 2024; https://github.com/shapely/shapely.

(23) Souza, P. C.; Alessandri, R.; Barnoud, J.; Thallmair, S.; Faustino, I.; Grünewald, F.; Patmanidis, I.; Abdizadeh, H.; Bruininks, B. M.; Wassenaar, T. A.; Kroon, P. C.; Melcr, J.; Nieto, V.; Corradi, V.; Khan, H. M.; Domański, J.; Javanainen, M.; Martinez-Seara, H.; Reuter, N.; Best, R. B.; Vattulainen, I.; Monticelli, L.; Periole, X.; Tieleman, D. P.; de Vries, A. H.; Marrink, S. J. Martini 3: a general purpose force field for coarse-grained molecular dynamics. Nature Methods 2021, 18, 382–388.

(24) Meng, E. C.; Goddard, T. D.; Pettersen, E. F.; Couch, G. S.; Pearson, Z. J.; Morris, J. H.; Ferrin, T. E. UCSF ChimeraX: Tools for structure building and analysis. Protein Science 2023, 32, e4792.

(25) Ingólfsson, H. I.; Bhatia, H.; Zeppelin, T.; Bennett, W. F. D.; Carpenter, K. A.; Hsu, P.-C.; Dharuman, G.; Bremer, P.-T.; Schiøtt, B.; Lightstone, F. C.; Carpenter, T. S. Capturing Biologically Complex Tissue-Specific Membranes at Different Levels of Compositional Complexity. The Journal of Physical Chemistry B 2020, 124, 7819–7829, PMID: 32790367.

(26) Hammond, A. T.; Heberle, F. A.; Baumgart, T.; Holowka, D.; Baird, B.; Feigenson, G. W. Crosslinking a lipid raft component triggers liquid ordered-liquid disordered phase separation in model plasma membranes. Proceedings of the National Academy of Sciences 2005, 102, 6320–6325.

(27) Hunter, J. C.; Gurbani, D.; Ficarro, S. B.; Carrasco, M. A.; Lim, S. M.; Choi, H. G.; Xie, T.; Marto, J. A.; Chen, Z.; Gray, N. S.; Westover, K. D. In situ selectivity profiling and crystal structure of SML-8-73-1, an active site inhibitor of oncogenic K-Ras G12C. Proceedings of the National Academy of Sciences 2014, 111, 8895–8900.

(28) Marcink, T. C.; Simoncic, J. A.; An, B.; Knapinska, A. M.; Fulcher, Y. G.; Akkaladevi, N.; Fields, G. B.; Van Doren, S. R. MT1-MMP Binds Membranes by Opposite Tips of Its Propeller to Position It for Pericellular Proteolysis. Structure 2019, 27, 281–292.e6.

(29) Cuello, L. G.; Jogini, V.; Cortes, D. M.; Perozo, E. Structural mechanism of C-type inactivation in K+ channels. Nature 2010, 466.

(30) Kutzner, C.; Köpfer, D. A.; Machtens, J.-P.; de Groot, B. L.; Song, C.; Zachariae, U. Insights into the function of ion channels by computational electrophysiology simulations. Biochimica et Biophysica Acta (BBA) - Biomembranes 2016, 1858, 1741–1752, New approaches for bridging computation and experiment on membrane proteins.

(31) Su, R.; Zeng, J.; Marcink, T. C.; Porotto, M.; Moscona, A.; O’Shaughnessy, B. Host Cell Membrane Capture by the SARS-CoV-2 Spike Protein Fusion Intermediate. ACS Central Science 2023, 9, 1213–1228.

(32) Saud, Z.; Tyrrell, V. J.; Zaragkoulias, A.; Protty, M. B.; Statkute, E.; Rubina, A.; Bentley, K.; White, D. A.; Rodrigues, P. D. S.; Murphy, R. C.; Köfeler, H.; Griffiths, W. J.; Alvarez-Jarreta, J.; Brown, R. W.; Newcombe, R. G.; Heyman, J.; Pritchard, M.; Mcleod, R. W.; Arya, A.; Lynch, C.-A.; Owens, D.; Jenkins, P. V.; Buurma, N. J.; O’Donnell, V. B.; Thomas, D. W.; Stanton, R. J. The SARS-CoV2 envelope differs from host cells, exposes procoagulant lipids, and is disrupted in vivo by oral rinses. Journal of Lipid Research 2022, 63, 100208.

(33) Klein, F.; Soñora, M.; Santos, L. H.; Frigini, E. N.; Ballesteros-Casallas, A.; Machado, M. R.; Pantano, S. The SIRAH force field: A suite for simulations of complex biological systems at the coarse-grained and multiscale levels. Journal of Structural Biology 2023, 215, 107985.

(34) Ingólfsson, H. I.; Melo, M. N.; van Eerden, F. J.; Arnarez, C.; Lopez, C. A.; Wassenaar, T. A.; Periole, X.; de Vries, A. H.; Tieleman, D. P.; Marrink, S. J. Lipid Organization of the Plasma Membrane. Journal of the American Chemical Society 2014, 136, 14554–14559.

(35) Ingólfsson, H. I.; Carpenter, T. S.; Bhatia, H.; Bremer, P. T.; Marrink, S. J.; Lightstone, F. C. Computational Lipidomics of the Neuronal Plasma Membrane. Biophysical Journal 2017, 113.

(36) Wassenaar, T. A.; Pluhackova, K.; Moussatova, A.; Sengupta, D.; Marrink, S. J.; Tieleman, D. P.; Böckmann, R. A. High-throughput simulations of dimer and trimer assembly of membrane proteins. The DAFT approach. Journal of Chemical Theory and Computation 2015, 11.

(37) Hilpert, C.; Beranger, L.; Souza, P. C.; Vainikka, P. A.; Nieto, V.; Marrink, S. J.; Monticelli, L.; Launay, G. Facilitating CG Simulations with MAD: The MArtini Database Server. Journal of Chemical Information and Modeling 2023, 63.

(38) Quemener, E.; Corvellec, M. SIDUS—the solution for extreme deduplication of an operating system. Linux J. 2013, 2013.

